# *Jagged1/Notch2* Controls Kidney Fibrosis via *Tfam*-mediated Metabolic Reprogramming

**DOI:** 10.1101/285726

**Authors:** Shizheng Huang, Jihwan Park, Chengxiang Qiu, Yasemin Sirin, Seung Hyeok Han, Szu-yuan Li, Verdon Taylor, Ursula Zimber-Strobl, Katalin Susztak

**Author notes:** Correspondence: Katalin Susztak.

## Abstract

While Notch signaling has been proposed to play a key role in fibrosis, the direct molecular pathways targeted by Notch signaling and the precise ligand and receptor pair that are responsible for kidney disease remain poorly defined.

In this study, we found that *JAG1* and *NOTCH2* showed the strongest correlation with the degree of interstitial fibrosis in a genome wide expression analysis of a large cohort of human kidney samples. RNA sequencing analysis of kidneys of mice with folic acid nephropathy, unilateral ureteral obstruction, or APOL1-associated kidney disease indicated that *Jag1* and *Notch2* levels were higher in all analyzed kidney fibrosis models. Mice with tubule-specific deletion of *Jag1* or *Notch2* (*Ksp^cre^/Jag1^flox/flox^*, and *Ksp^cre^/Notch2^flox/flox^*) had no kidney-specific alterations at baseline, but showed protection from folic acid induced kidney fibrosis. Tubule-specific genetic deletion of *Notch1* and global knock-out of *Notch3* had no effect on fibrosis. *In vitro* chromatin immunoprecipitation experiments and genome-wide expression studies identified the mitochondrial transcription factor A (*Tfam*) as a direct Notch target. Re-expression of *Tfam* in tubule cells prevented Notch-induced metabolic and profibrotic reprogramming. Kidney tubule specific deletion of *Tfam* resulted in perinatal lethality.

In summary, *Jag1/Notch2* plays a key role in kidney fibrosis development by regulating *Tfam* expression and metabolic reprogramming.

## Introduction

One in ten people worldwide suffers from chronic kidney disease (CKD) (1, 2). Fibrosis is the final common pathway and histological manifestation of CKD which is characterized by the accumulation of collagen, activated myofibroblasts, inflammatory cells, epithelial cell dedifferentiation, and loss of vascular supply in the kidney (3–5).

The mechanism of fibrosis is a hotly debated issue. Renal tubule epithelial cells (RTEC) appear to play key role in fibrosis development. Direct *in vivo* genetic manipulations of RTEC is both sufficient to induce or to ameliorate fibrosis development in mice (6–8). Injury to RTECs is usually agreed to be a proximal lesion in the downstream cascade that eventually results in fibrosis (9, 10). Recent studies highlighted the reemergence of developmental pathways in fibrosis including Notch, Wnt and Hedgehog in response to severe tubule injury (11–16). While chronic activation of Notch plays a key role in development and regeneration of the kidney, sustained (tonic) expression of Notch in RTEC will expand the progenitor (transit) amplifying pool but eventually block full differentiation of RTEC. Epithelial dedifferentiation, due to the lack of expression of key transporter and functional proteins will lead to CKD development. Direct targets of Notch signaling and the exact mechanism of Notch-induced RTEC dedifferentiation is not fully understood.

Notch signaling is a highly conserved cell-cell communication mechanism that regulates tissue development, homeostasis, and repair. In mammals, there are four Notch receptors (*NOTCH 1-4*) and five canonical ligands, Jagged (*JAG1, 2*) and Delta-like ligand (*DLL1, 3, 4*). Upon ligand binding, the Notch intracellular domain (NICD) travels to the nucleus, binds to RBPJ (recombination signal binding protein for immunoglobulin kappa J region), and mediates gene transcription (17). While Notch signaling plays an important role in a multitude of disease development, it also has an important homeostatic function (16). Systemic pharmacological targeting of NOTCH has been developed and is in clinical trials for various solid cancers (18). Most of these drugs have been associated with significant toxicities that which likely limit their development for chronic, slowly progressing conditions such as fibrosis (18–21). New results on the other hand suggest that while different Notch ligands and receptors are homologous, they seem to exert isoform-specific functions (16). By using antibodies to specifically target Notch signaling, the Jones’ group found that inhibition of JAG1, acting together with NOTCH2, reduces tumor burden in a mouse model of primary liver cancer (22). Tran *et al* demonstrated the dominant role of DLL4-NOTCH1 signaling in T-cells during graft-versus-host disease using a ligand-targeted antibody strategy (23). Ligand-specific targeting strategies were able to dissociate disease causing and homeostatic functions of Notch signaling; therefore, they could offer novel therapeutic strategies.

Our previous results indicated that increased and sustained tubular epithelial Notch signaling following injury plays a key role in the development of the renal tubulointerstitial fibrosis (TIF) (8, 24–28). Global and systemic inhibition of Notch signaling using small molecular blockers of the gamma secretase pathway ameliorated fibrosis in the folic acid, unilateral ureteral obstruction, and HIV-induced kidney fibrosis models (29–34). Similarly, mice with genetic deletion of *Rbpj* in renal tubule cells were protected from kidney fibrosis development. As RBPJ is a common partner for all Notch receptors, these studies were unable to identify the specific Notch ligand/receptor pair responsible for the profibrotic effect of Notch signaling. Furthermore, RBPJ also has Notch independent activity. Identification of specific ligands and receptors for kidney fibrosis therefore remains a critically important issue.

Here, we analyzed several mouse models and patient samples with kidney disease to understand the expression of Notch ligands and receptors. We generated mouse models with genetic deletion of individual Notch ligands and receptors, and finally, we identified direct targets of Notch signaling. We show that RTEC expression of *Jag1* acting together with *Notch2* leads to metabolic reprogramming of RTEC via *Tfam*, resulting in epithelial dedifferentiation and fibrosis development.

## Results

### Increased *Jag1* and *Notch2* expression in RTEC of mice and patients with kidney fibrosis

To identify Notch ligands and receptors responsible for fibrosis development, we first evaluated results from unbiased gene expression (RNA sequencing) studies of mice with kidney fibrosis, including folic acid (FA) nephropathy, unilateral ureteral obstruction (UUO) model, and podocyte-specific expression of APOL1 risk variants (Figure 1, **A-C**) (35–37). Amongst the different ligands, *Jag1* transcript level was the highest in whole kidney lysates while the level of *Dll3* was too low to detect. Expression of *Jag1* was consistently and significantly increased in kidneys of all examined CKD models (Figure 1, **A-C**). Of the Notch receptors, we found *Notch2* had the highest expression in the kidney and its level was consistently increased in all fibrosis models.

**Figure 1.**
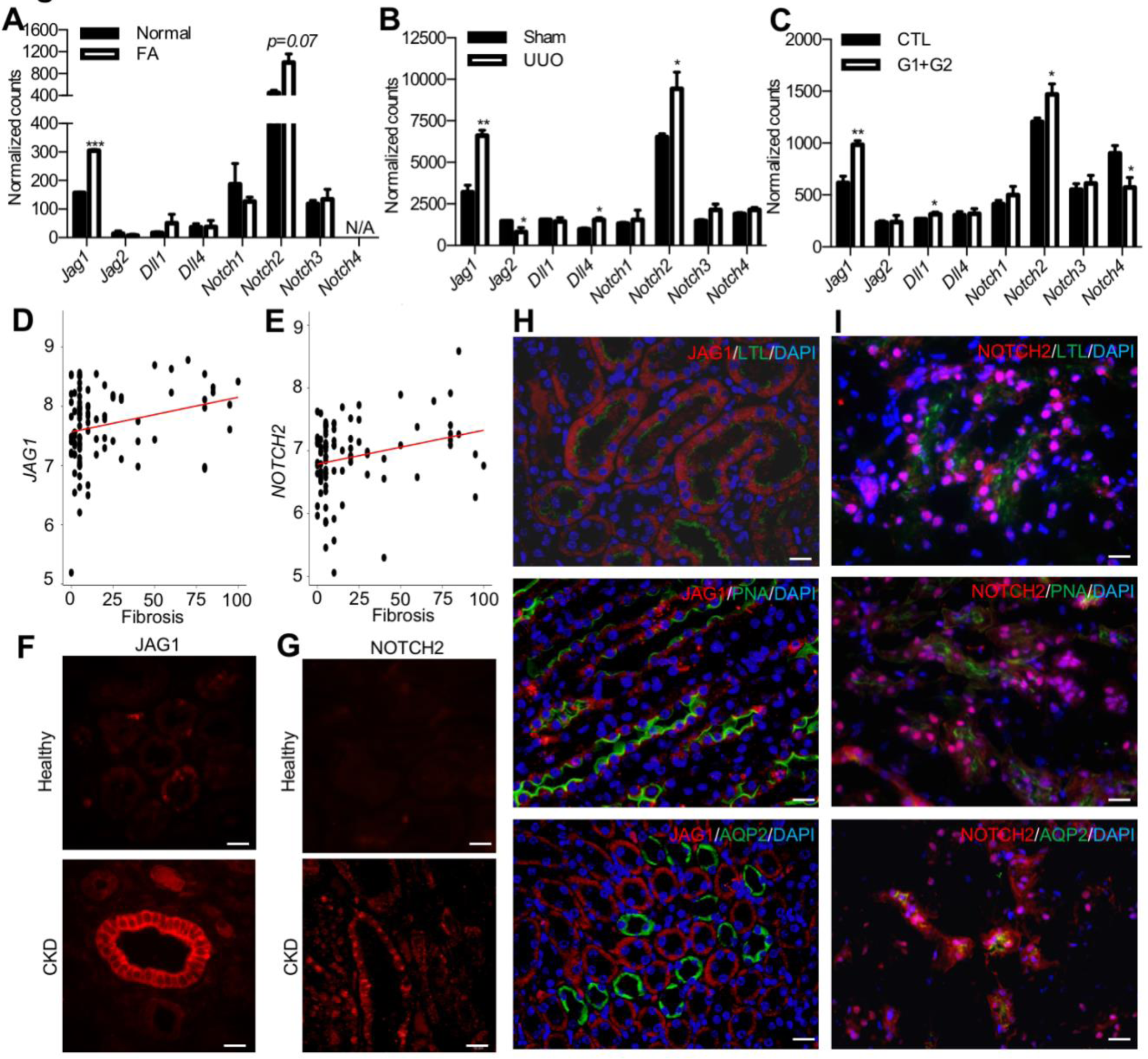
Increased *Jag1/Notch2* expression in mouse and human kidney fibrosis. **(A-C)** Normalized counts of Notch ligands and receptors by RNA sequencing in whole-kidney lysates of folic acid-induced nephropathy group (A), sham and UUO group (B), and APOL1-G1/G2 mice (C). Data are represented as the mean ± SEM. \**P* < 0.05, ** *P* < 0.01, *** *P* < 0.001, by 2-tailed Student’s *t* test, *n*=2-6 per group. **(D and E)** Correlation between interstitial fibrosis and *JAG1* (D) or *NOTCH2* (E) transcript level in 95 microdissected human kidney samples. **(F and G)** Representative images of immunofluorescence staining with antibodies against JAG1 (F) and NOTCH2 (G) in healthy and diseased human samples. Scale bar: 10µm. **(H and I)** Double staining of JAG1 (H) and NOTCH2 (I) with LTL, PNA, or AQP2 on FA treated mouse kidneys. Scale bar: 10µm.

To understand the human translatability of the studied mouse models, we have analyzed gene expression changes in 95 microdissected human kidney tubule samples. The dataset included microarray samples from healthy, diabetic, and hypertensive subjects, and patients with varying degrees of fibrosis and chronic kidney disease (38). Among all tested Notch ligands, *JAG1* expression showed significant positive correlation with the degree of fibrosis (*P=0.003*) (Table 1; Figure 1D). We also found a positive association between the expression of *NOTCH2* with the degree of interstitial fibrosis (*P=0.02*) (Figure 1E).

**Table 1:** Correlation between interstitial fibrosis and Notch signaling transcript level in 95 microdissected human kidney samples.

|  | Beta Value | *P* Value |
| --- | --- | --- |
| *JAG1* | 0.0078 | 0.0039 |
| *JAG2* | 0.0031 | 0.0889 |
| *NOTCH1* | -0.0001 | 0.9313 |
| *NOTCH2* | 0.0062 | 0.0215 |
| *NOTCH3* | 0.0029 | 0.2675 |
| *NOTCH4* | 0.0039 | 0.1085 |

Given that Notch is activated by proteolytic cleavage, we then confirmed the increase in cleaved protein expression of JAG1 and NOTCH2 in kidneys with fibrosis. Immunostaining studies indicated increased protein expression of JAG1 in kidney tubule of human TIF samples. Only few cells were positive for JAG1 in healthy human kidney samples, but its expression was much increased in RTEC of kidneys obtained from CKD subjects (Figure 1F). Similar results were obtained for NOTCH2 (Figure 1G).

We next examined JAG1 and NOTCH2 level in kidneys of FA-induced kidney fibrosis mice.

Double immunostaining of JAG1 with kidney-segment-specific markers identified JAG1-positive cells in LTL (lotus lectin, a marker for proximal tubule) and PNA (peanut agglutinin, a marker for distal tubule/loop of Henle) positive tubules, but not in AQP2 (aquaporin 2, a marker for collecting duct) expressing tubule segments. We also identified NOTCH2-positive cells in LTL-, PNA- and AQP2-positive tubule segments, indicating that most examined tubule segments expressed NOTCH2 (Figure 1, **H and I**). In summary, *Jag1* and *Notch2* were the highest expressed Notch ligands and receptors in whole kidney lysates, and both their transcript and protein levels showed a consistent increase in kidney tubule cells in patient samples and in animal models.

### TIF development is mediated by tubule epithelial expression of *Jag1* and *Notch2*

Next, we studied whether increased JAG1 expression observed in CKD renal tubule cells contributes to fibrosis development. Therefore, we crossed the *Jag1^flox/flox^* animals with mice expressing Cre recombinase under the cadherin 16 promoter to generate animals with *Jag1* deletion in RTEC (*Ksp^cre^/Jag1^flox/flox^*) (Figure 2A) (39). *Ksp^cre^/Jag1^flox/flox^* mice showed lower *Jag1* expression but no significant histological alterations at baseline (Figure 2B). Kidney fibrosis was induced by FA injection. We found that 7 days after FA injection, kidney histology was markedly improved in the *Ksp^cre^/Jag1^flox/flox^* mice when compared to control animals as it was evident by Periodic acid-Schiff (PAS)-stained kidney sections (Figure 2B). Serum blood urea nitrogen (BUN) and creatinine indicated functional improvement in tubule-specific *Jag1* knockout mice (Figure 2C **and D**). There was marked reduction in the expression of *Notch1-3* and Notch target genes including *HeyL* in FA-treated *Ksp^cre^/Jag1^flox/flox^* mice, indicating the successful deletion of Notch signaling. Epithelial dedifferentiation and proliferation are the hallmarks of fibrosis. Expression levels of *Fibronectin, Vimentin, Collagen1, 3, Snai1, Snai2* and *c-Myc* were lower in the folic acid injected *Jag1* knock-out mice when compared to littermate controls (Figure 2E **and F**).

**Figure 2:**
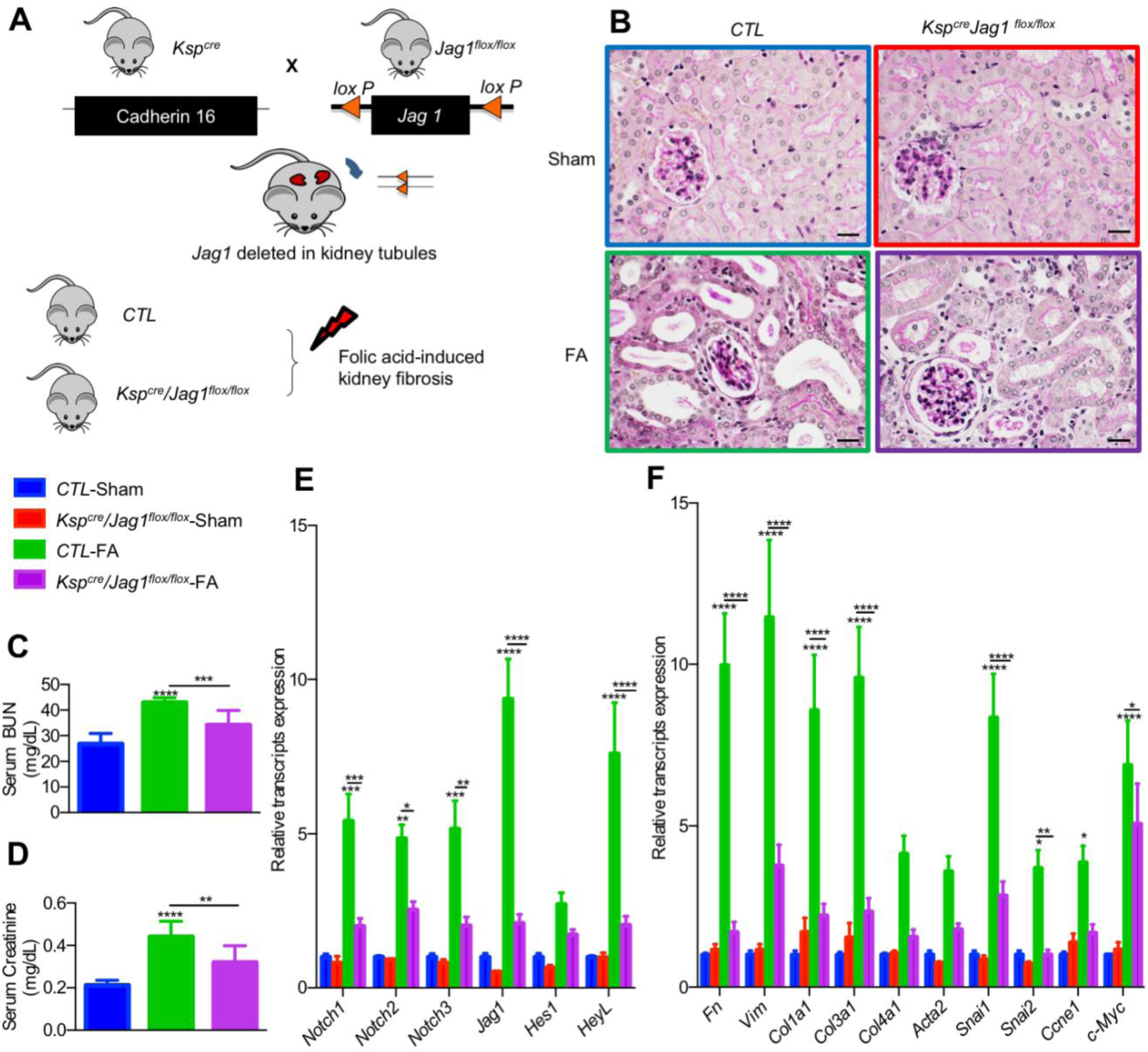
Reduced fibrosis in tubule specific *Jag1* knock-out mice. **(A)** Experimental scheme for generating the *Ksp^cre^ Jag1^flox/flox^* mice. Kidney injury was induced by FA injection. **(B)** Representative images of PAS-stained kidney sections from control and *Ksp^cre^ Jag1^flox/flox^* mice with or without FA injection. Scale bar: 10µm. (**C and D**) Serum BUN and creatinine measurement of control and *Ksp^cre^ Jag1^flox/flox^* mice with or without FA injection. Data are represented as mean ± SEM. ** *P* <0.01, *** *P* <0.001, **** *P* < 0.0001, by One-way ANOVA with *post-hoc* Tukey test, *n*=5-10 per group. **(E and F)** Relative mRNA amount of Notch signaling (E), fibrosis markers, dedifferentiation and proliferation (F) in control and *Ksp^cre^ Jag1^flox/flox^* mice with or without FA injection. Data are represented as mean ± SEM. * *P* <0.05; ** *P* <0.01, *** *P* <0.001, **** *P* < 0.0001, by Two-way ANOVA with *post-hoc* Tukey test, *n*=3-10 per group.

Our unbiased analysis identified *NOTCH2* as a Notch receptor with the strongest correlation with fibrosis. We next studied the role of *Notch2* in kidney fibrosis. We generated mice with RTEC-specific deletion of *Notch2* by mating the *Notch2^flox/flox^* mice with the *Ksp*^cre^ mice (Figure 3A). Animals were born with expected Mendelian frequency and histological analysis showed no significant difference between control and conditional knock-out mice (Figure 3B). Folic acid injection was used to induce kidney fibrosis. The expression of *Notch2 Notch3* and *HeyL* levels were significantly decreased while *Notch1* and *Jag1* levels were unchanged in FA-treated *Ksp*^*cre*^*Notch2*^*flox/flox*^ mice (Figure 3C). We found marked structural improvement when PAS-stained kidney sections of folic acid injected control and *Ksp*^*cre*^*Notch2*^*flox/flox*^ were compared (Figure 3B). In conjunction, levels of the profibrotic genes *Fibronection, Vimentin, Collagen1, 3, Acta2, Snai1 and Snai2* were significantly reduced in tubule-specific *Notch2* knock-out animals (Figure 3D).

**Figure 3:**
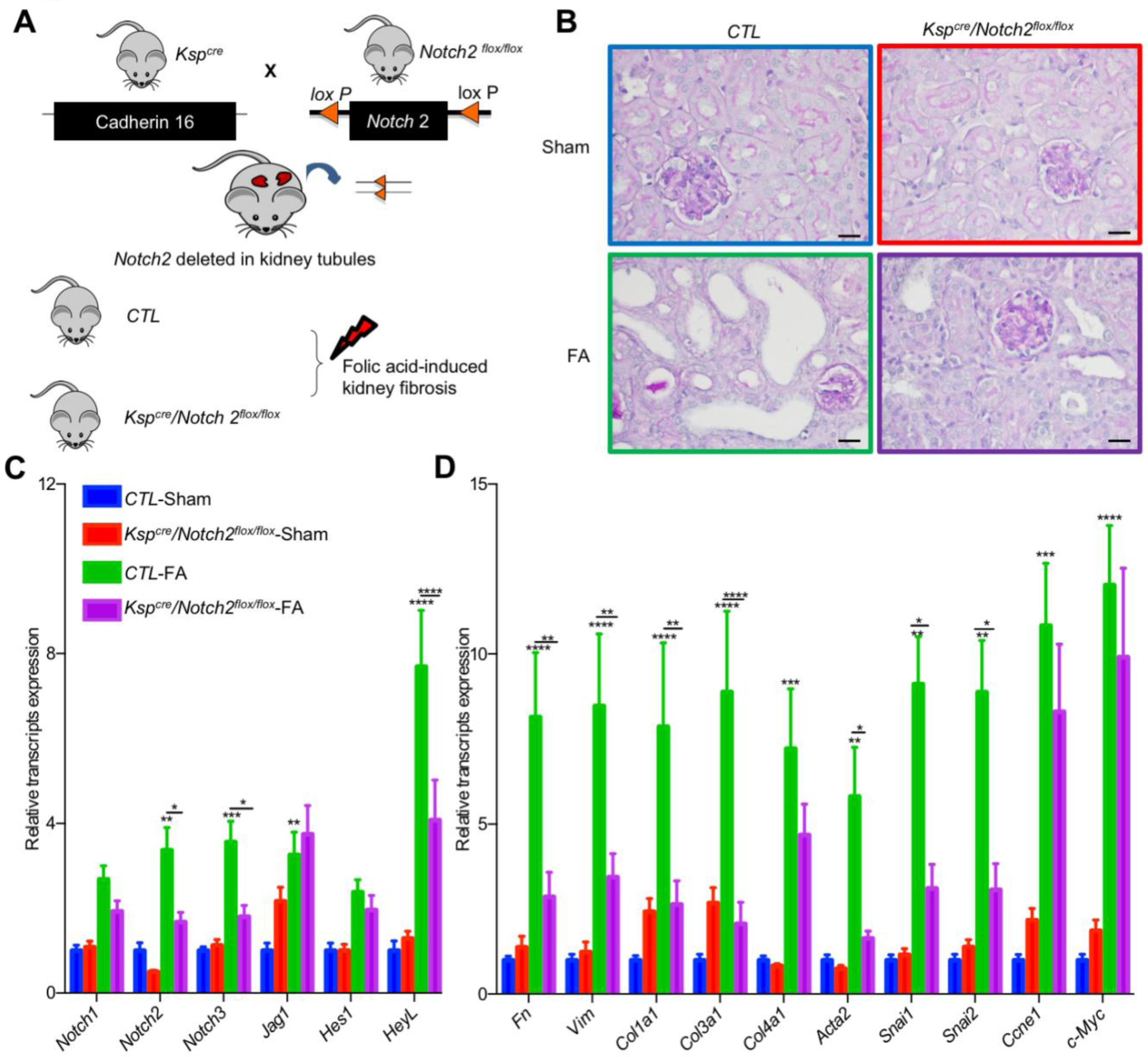
Reduced fibrosis in tubule specific *Notch2* knock-out mice. **(A)** Experimental scheme for generating the *Ksp^cre^ Notch2 ^flox/flox^* mice. Kidney injury was induced by FA injection. **(B)** Representative images of PAS-stained kidney sections from control and *Ksp^cre^ Notch2^flox/flox^* mice with or without FA injection. Scale bar: 10µm. **(C and D)** Relative mRNA amount of Notch signaling (C), fibrosis, dedifferentiation and proliferation markers (D) in control and *Ksp^cre^ Notch2^flox/flox^* mice with or without FA injection. Data are represented as mean ± SEM. * *P* <0.05; ** *P* <0.01, *** *P* <0.001, **** *P* < 0.0001, by Two-way ANOVA with *post-hoc* Tukey test, *n*=5-7 per group.

To understand whether this effect is specific for *Jag1* and *Notch2*, we tested the effect of genetic deletion of *Notch1* and *Notch3*. Tubule-specific *Notch1* knock-out mice was generated by mating the *Ksp^cre^* and *Notch1*^flox/flox^ animals. We found no significant differences in kidney fibrosis development when the *Ksp*^*cre*^*Notch1*^flox/flox^ were compared to control FA-treated animals (**Supplemental Figure 1, A-D**). Similarly, we studied mice with global deletion of *Notch3*. We found no renal abnormalities in global *Notch3* knock-out mice at baseline and *Notch3* knockout mice showed no significant differences in FA-induced kidney fibrosis development (**Supplemental Figure 2, A-D)**. In summary, *in vivo* studies indicate that tubule specific deletion of *Jag1* and *Notch2* ameliorates kidney fibrosis development in mice while *Notch1* and *Notch3* minimally contribute to CKD.

### Jag1/Notch2 in RTEC induces dedifferentiation and proliferation

In order to understand the downstream molecular pathways regulated by *Jag1/Notch2*, we turned to an *in vitro* system. TGF-β1 is one of the most powerful profibrotic cytokines and a known inducer of Notch signaling in epithelial cells by *Smad3* (8, 40). TGF-β1 treatment significantly increased *Jag1* expression in primary culture of RTECs (Figure 4A), indicating that TGF-β1 is an upstream Notch regulator in tubule cells. Similar to the *in vivo* kidney data, the expression of *Notch2* was the highest amongst the different Notch receptors in cultured RTECs.

**Figure 4:**
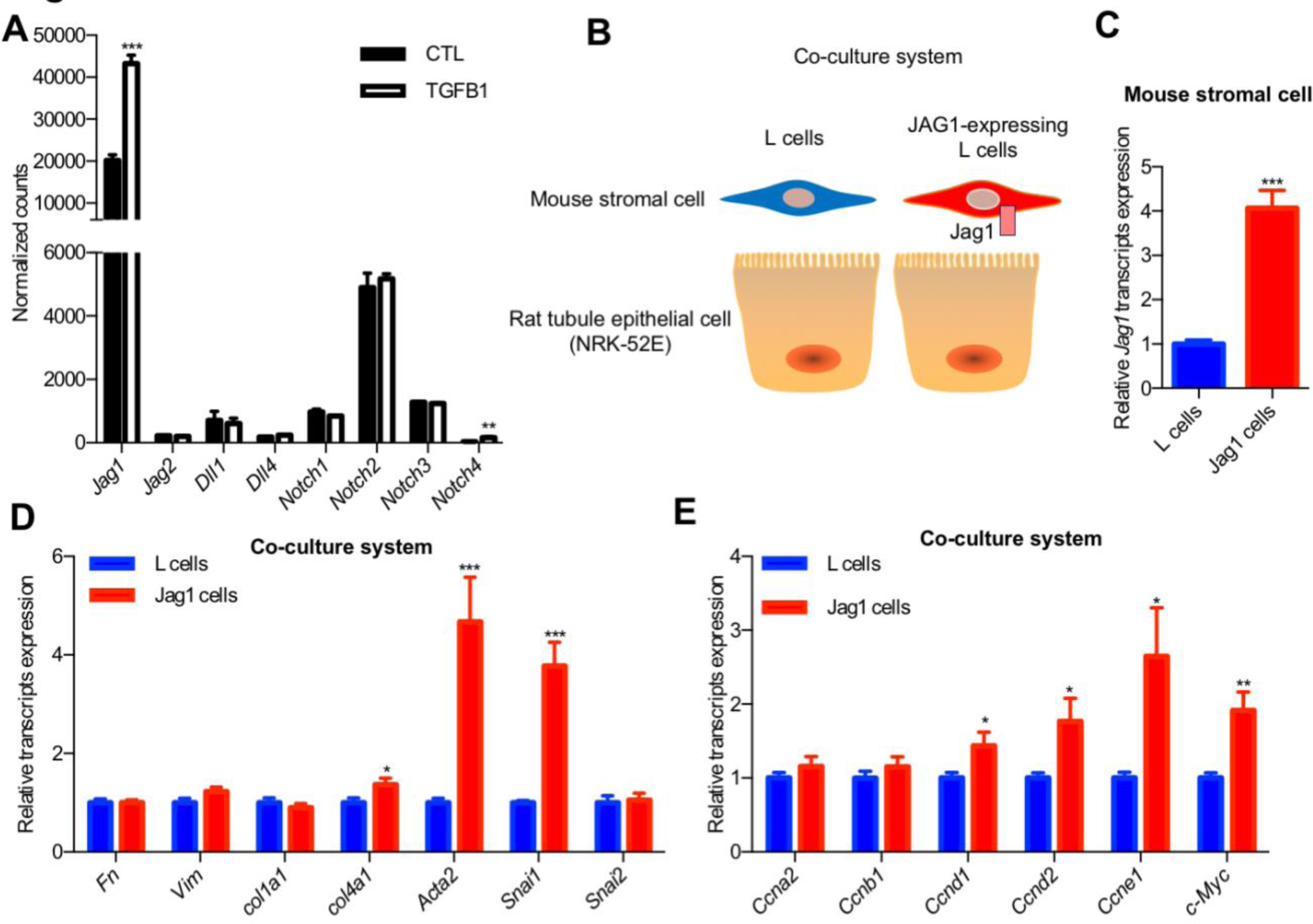
*Jag1* expression induces dedifferentiation and proliferation of RTEC. **(A)** Normalized counts of Notch ligands and receptors transcripts of TGFβ1-treated primary cultured mouse RTECs. Data are represented as the mean ± SEM. ** *P* < 0.01, *** *P* < 0.001, by 2-tailed Student’s *t* test, *n*=3 per group. **(B)** Experimental scheme for JAG1 co-culture system. **(C)** Expression of *Jag1* in JAG1-expressing L cell compared to control L cell. Data are represented as the mean ± SEM. *** *P* < 0.001, by 2-tailed Student’s *t* test, *n*=12 per group. **(D and E)** Co-culturing of JAG1-expressing cells with NRK-52E causing cell proliferation and dedifferentiation. * *P* < 0.05, \*\**P <* 0.01, *** *P* < 0.001, by 2-tailed Student’s *t* test, *n*=12 per group.

Furthermore, to study the role of JAG1 specifically, we have set-up a co-culture system by mixing RTEC with JAG1-expressing or control stromal cells (Figure 4, B and C). In this system, as the ligand expressing (mouse stromal cells) and signal receiving cells (rat kidney epithelial cells; NRK-52E) are from different species, species specific primers can differentiate the contribution of signal sending from signal receiving cells.

Culturing NRK-52E with JAG1-expressing stromal cells resulted in increase in expression of markers of dedifferentiation such as *Acta2*, *Snai1* and enhanced proliferation such as an increase in expression of *Cyclins* and *c-Myc*, when compared to control cells (Figure 4, D and E). The *in vitro* regulation of *Acta2*, *Snai1* and *c-Myc* by JAG1/NOTCH2 recapitulated the *in vivo* mouse model results (Figure 2F and Figure 3D). JAG1/NOTCH2 activation resulted in epithelial dedifferentiation and proliferation both *in vivo* in mouse models and *in vitro* in tubule cells.

### Tfam is direct target of Notch

To understand the direct molecular targets of Notch signaling, we have examined RBPJ chromatin immunoprecipitation datasets (ChIP-seq) (41). To filter out functionally important binding that are also associated with gene expression changes *in vivo*, we correlated the ChIP-seq results with changes observed by transcriptome analysis of kidneys from tubule-specific Notch1 transgenic mice (*Pax8^rtTA^/TRE^ICNotch1^*) (8). This combined analysis identified 79 RBPJ-binding sites and those gene expression also changed following *in vivo* NOTCH expression in tubule cells (56 increase and 23 decrease RBPJ-binding sites) (**Supplemental Figure 3A; Supplemental Table1 and 2**). Gene ontology (GO) classification of genes with direct RBPJ binding and RTEC gene expression changes indicated enrichment for metabolic pathways (i.e. *Rxra, Tfam, Acot12, Ndufa10*, etc) (Table 2).

**Table 2:**
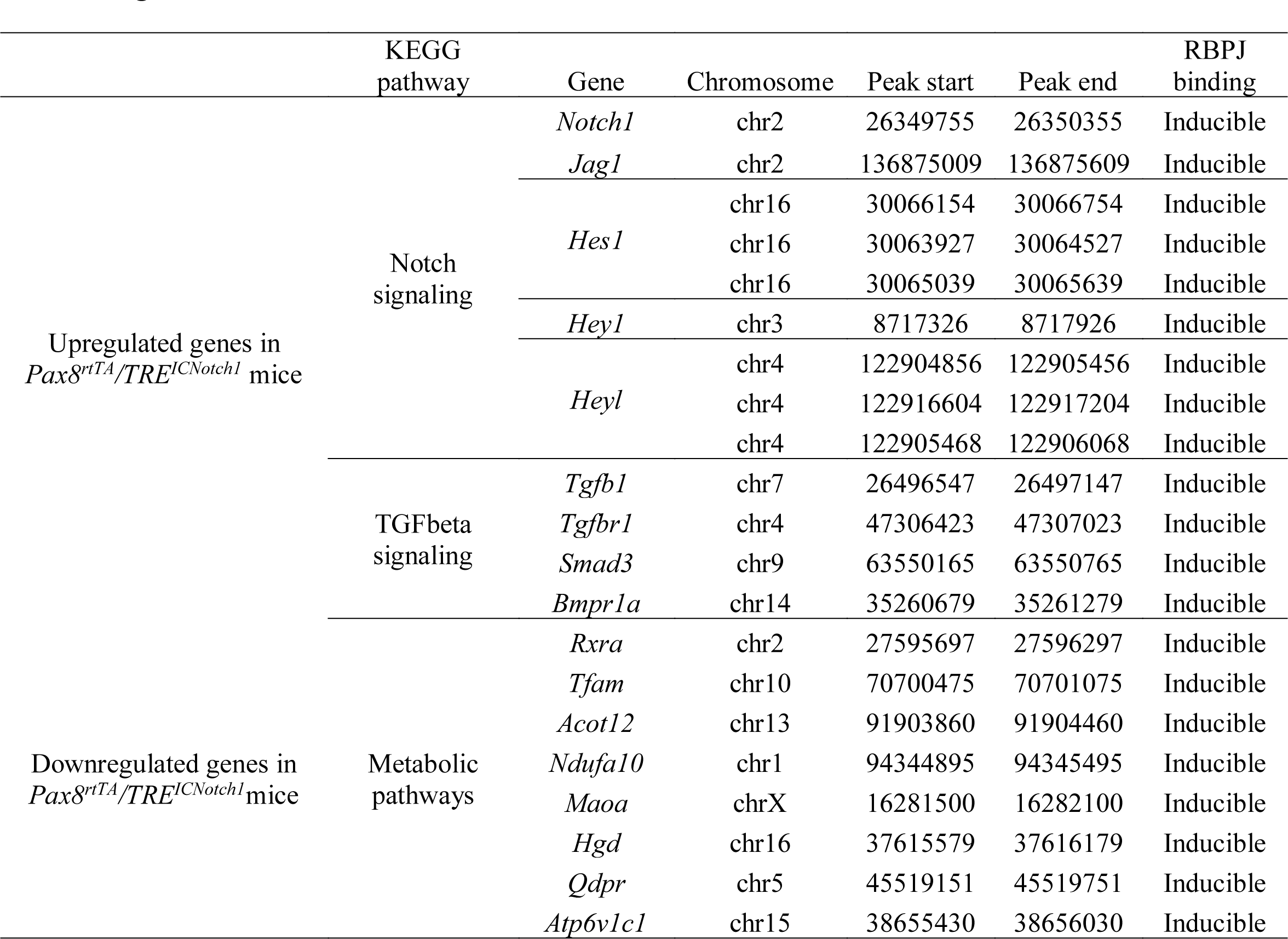
Gene ontology analysis of overlap RBPJ binding peaks identified in Supplemental Figure 3A.

Epidermal growth factor (EGF, encoded by *Egf*) was one of the top differentially expressed genes in *Pax8^rtTA^/TRE^ICNotch1^* mice (**Supplemental Table2**) with lower expression in Notch transgenic animals. We found that RBPJ could directly bind to the *Egf* regulatory region (**Supplemental Figure 4A**). *EGF* showed strong negative correlation with kidney fibrosis score in human kidney samples (*P*=8.91×10^−7^) (**Supplemental Figure 4B**). *Egf* was also persistent decreased in FA, UUO, and APOL1 CKD mouse models (**Supplemental Figure 4, C-E**). To understand the role of EGF, we added Recombinant Human EGF to culture medium. We found that exogenous EGF was unable to ameliorate Notch-induced expression of *Acta2* and *Snai1* (**Supplemental Figure 4F**). In summary, we were unable to connect the Notch-induced *Egf* changes to fibrosis development.

From the list of direct target genes, we next focused on *Tfam*. *Tfam* is one of the key transcriptional regulators of mitochondrial (and therefore metabolic) gene expression. ChIP-seq analysis indicated RBPJ binding to the *Tfam* regulatory region (at chr10: 70700475-70701075) (Figure 5A).

**Figure 5:**
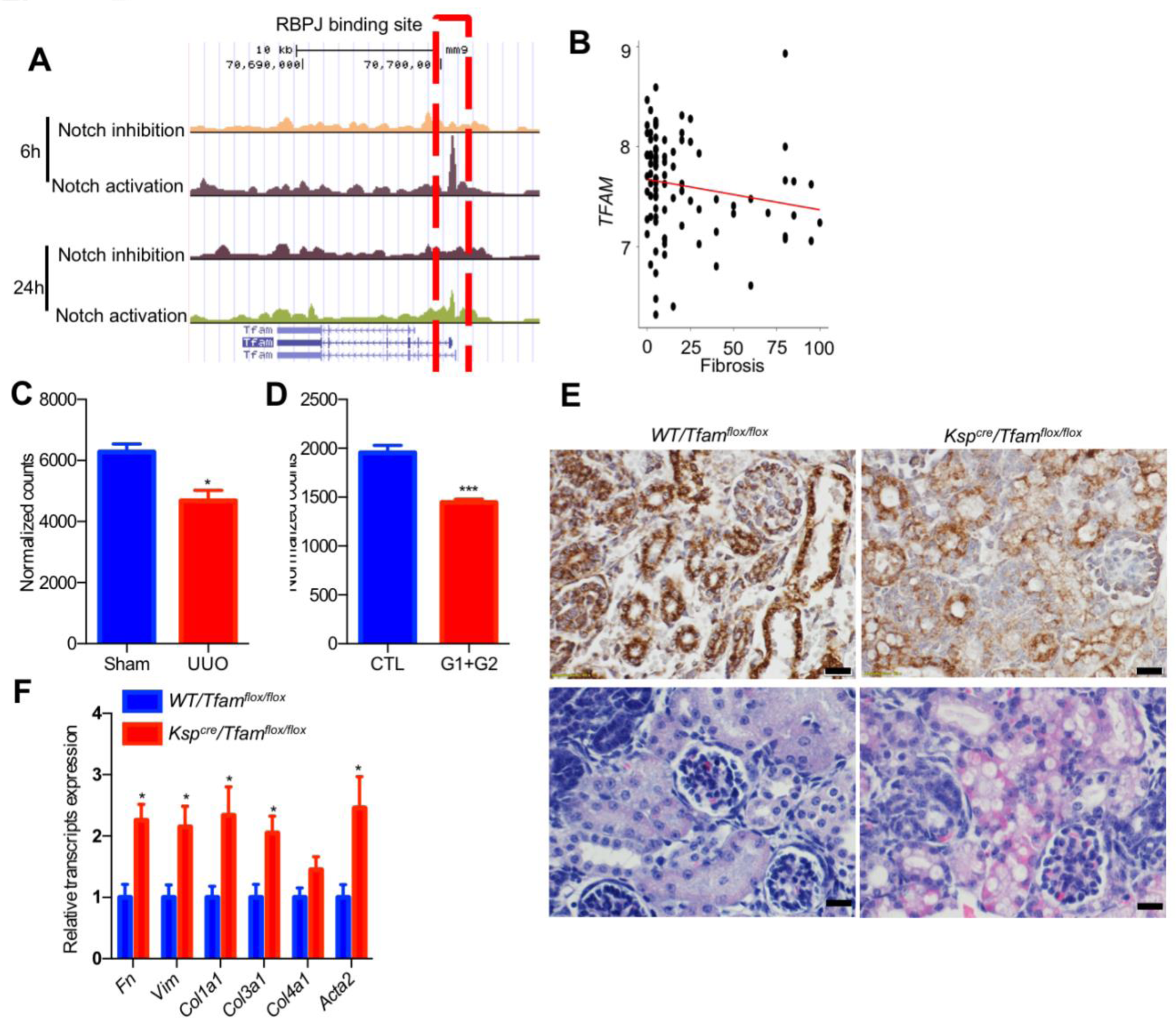
*Tfam* is the direct target of Notch and plays an important role in kidney tubules. **(A)** The mouse *Tfam* locus and RBPJ ChIP-Seq from 6h and 24h Notch activation and inhibition. **(B)** Correlation between interstitial fibrosis and *TFAM* transcript level in 95 microdissected human kidney samples. **(C and D)** Normalized counts of *Tfam* by RNA sequencing in whole-kidney lysates of sham and UUO group (C), and APOL1-G1/G2 mice (D). Data are represented as the mean ± SEM. * *P* < 0.05, *** *P* < 0.001, by 2-tailed Student’s *t* test, *n*=3-6 per group. **(E)** COX IV and H&E staining from kidneys of *Ksp^cre^Tfam^flox/flox^* mice and control littermates. Scale bar: 10µm. **(F)** Relative mRNA amount of fibrosis markers in control and *Ksp^cre^ Tfam^flox/flox^* mice. Data are represented as mean ± SEM. * *P* <0.05, by 2-tailed Student’s *t* test, *n*=3-4 per group.

*TFAM* was not only downregulated directly by Notch signaling, but its expression was also negatively associated with the degree of kidney fibrosis in human patient kidney samples (*P*=0.05) (Figure 5B). *Tfam* expression was decreased not only in tubule specific Notch transgenic mice, but also in the UUO and APOL1-induced kidney fibrosis models (Figure 5, C and D).

While previous studies indicated that peroxisome proliferator-activated receptor γ coactivator-1α (PGC1α, encoded by the gene *Ppargc1a*) can rescue Notch-induced kidney fibrosis, the reduced PGC1α cannot be responsible for Notch-induced cellular dedifferentiation as the PGC1α knock-out mice has no renal phenotype at baseline(42, 43). To understand whether reduced *Tfam* expression plays a role in tubule epithelial dedifferentiation, we generated a new mouse model with tubule-epithelial-specific deletion of *Tfam* by mating the *Tfam^flox/flox^* and *Ksp^Cre^* mice (**Supplemental Figure 3B**). We found fewer than expected *Ksp^Cre^Tfam^flox/flox^* mice in the litter when compared to heterozygous or wild type controls (Table 3). Most *Tfam* knockout mice died shortly after birth, and the few surviving animals were significantly smaller by postnatal day 15 than control littermates (**Supplemental Figure 3C**). Histological examination indicated mitochondrial alterations in tubule epithelial cells of *Ksp^Cre^Tfam^flox/flox^* mice, indicating the key role of Tfam in renal tubule homeostasis and metabolism (Figure 5E). We found that *Ksp^Cre^Tfam^flox/flox^* mice had profibrotic gene expression changes as indicated by higher levels of *Fibronectin, Vimentin, Collagen1, 3* and *Acta2* on postnatal day 20 (Figure 5F). Overall, we found a direct binding of RBPJ into the *Tfam* promoter and reduced *Tfam* expression following Notch activation. *Tfam* expression in tubule cells were critically important for kidney function as mice with genetic deletion of *Tfam* in tubule cells died shortly after birth.

**Table 3:**
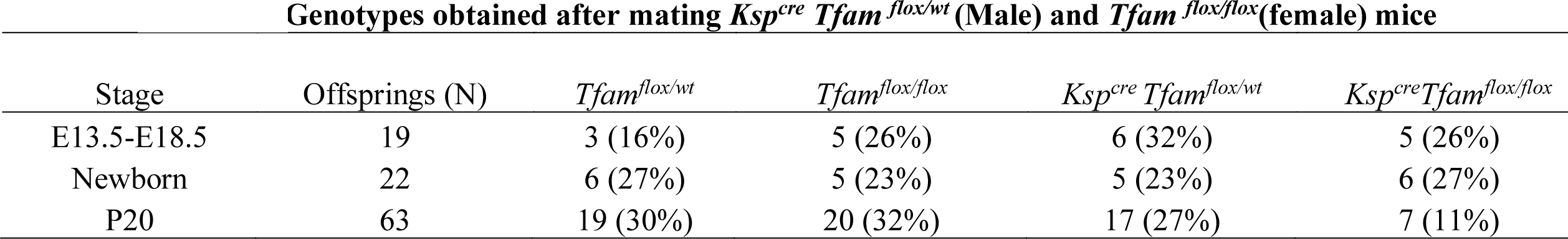
Genotypes obtained after mating *Ksp^cre^ Tfam ^flox/wt^* (Male) and *Tfam ^flox/flox^*(female) mice

### Tfam can rescue Notch-induced metabolic defect and downstream profibrotic changes

As we identified *Tfam* as a direct Notch target, we hypothesized that the widespread effect of Notch on gene expression and RTEC dedifferentiation is mediated by direct metabolic reprogramming. To understand the functional consequences of Notch-induced reduction of *Tfam* expression, we have examined mitochondrial function in RTEC. RTEC exclusively rely on mitochondrial fatty acid oxidation (FAO) as their energy source (44, 45). First, we examined *Tfam* expression *in vitro* following *Jag1/Notch2* activation. *Tfam* expression was lower in RTEC-cultured in cells treated with TGF-β1 and also in the presence of JAG1-expressing stromal cells (Figure 6A; **Supplemental Figure 5A**). We found that increased Notch activation was associated with a decrease in FAO of tubule cells confirming that *Tfam* is a direct Notch target. Increasing *Tfam* expression in RTEC improved the FAO defect both in the TGF-β1-induced Notch activation models and in the JAG1 co-culture system (Figure 6, **B and C; Supplemental Figure 5, B and C**), thus indicating that *Tfam* is functionally important in mediating the Notch-induced metabolic reprogramming.

**Figure 6:**
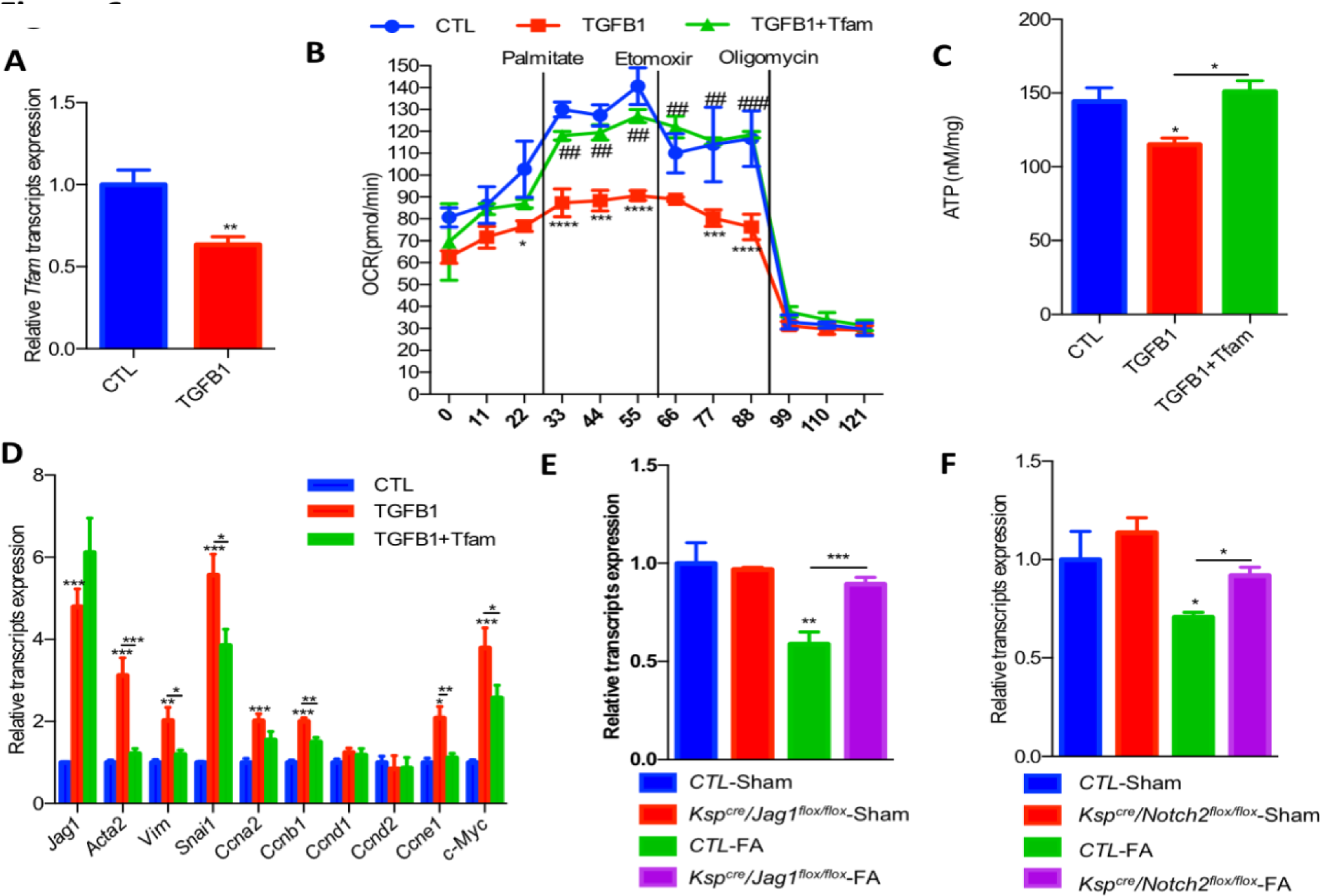
*Tfam* rescues TGF-β1-induced metabolic defect and downstream profibrotic changes. **(A)** *Tfam* expression level in NRK-52E cells treated with TGF-β1 for 24h. Data are represented as mean ± SEM. ** *P* <0.01, by 2-tailed Student’s *t* test, *n*=8 per group. **(B)** Oxygen consumption rate (OCR) in NRK-52E cells exposed to 5 ng/ml TGF-β1 for 24 h in the presence of GFP or *Tfam* plasmid. Where indicated, cells were incubated in palmitate (180 μM), etomoxir (40μM) and oligomycin (1 μM). Data are represented as mean ± SEM. \**P* < 0.05, *** *P* <0.001 and **** *P* <0.0001 as compared to control (CTL) group, ##*P* < 0.01, ### *P* <0.001 as compared to TGF-β1 group, by Two-way ANOVA with *post-hoc* Tukey test, *n*=3 per group. (**C**) ATP levels in NRK52E cells and cells treated with TGF-β1 for 24h in the presence of GFP or *Tfam* plasmid. Data are represented as mean ± SEM. * *P* <0.05, by One-way ANOVA with *post-hoc* Tukey test, *n*=7-8 per group. **(D)** Relative mRNA expression of transcripts related to fibrosis, dedifferentiation and proliferation in NRK52E cells treated with or without TGF-β1 and GFP or *Tfam* plasmid. Data are represented as mean ± SEM. \**P* < 0.05, \*\**P* < 0.01, \*\*\**P* < 0.001 by One-way ANOVA with Tukey’s *post hoc* tests, *n*=7-8 per group. (**E**) Relative mRNA amount of *Tfam* in control and *Ksp^cre^Jag1^flox/flox^* mice with or without FA injection. Data are represented as mean ± SEM. **, *P* <0.01; ***, *P* <0.001, by Two-way ANOVA with *post-hoc* Tukey test, *n*=3-10 per group. (**F**) Relative mRNA amount of *Tfam* in control and *Ksp^cre^Notch2^flox/flox^* mice with or without FA injection. Data are represented as mean ± SEM. *, *P* <0.05, by Two-way ANOVA with *post-hoc* Tukey test, *n*=5-7 per group.

Furthermore, the improved FAO and mitochondrial content significantly ameliorated the Notch induced profibrotic gene expression changes (Figure 6D; **Supplemental Figure 5D**). To prove that the rescue effect was related to improved FAO, we have also treated cells with the peroxisome proliferator-activated receptor α (PPARα) agonist, fenofibrate. We found that fenofibrate could rescue Jag1-induced fibrosis (**Supplemental Figure 6, A and B**), indicating the key role of metabolic changes in Notch induced fibrosis.

Finally, we have examined whether *Jag1* and *Notch2* deletion can rescue from the metabolic defect observed in mouse kidney fibrosis samples. We found that *Tfam* expression was decreased in the FA-induced kidney fibrosis in control groups. Genetic deletion of *Jag1* and *Notch2* prevented the decrease in *Tfam* expression and downstream fibrosis development (Figure 6, E and F). On the other hand, mice with genetic deletion of *Notch1* and *Notch3* were not protected from fibrosis induced a decrease in *Tfam* expression **(Supplemental Figure 5, E and F**). In summary, these results strongly indicate that the Notch-induced inhibition of *Tfam* and downstream metabolic reprogramming are responsible for the profibrotic effect of Notch both *in vivo* and *in vitro* (Figure 7).

**Figure 7:**
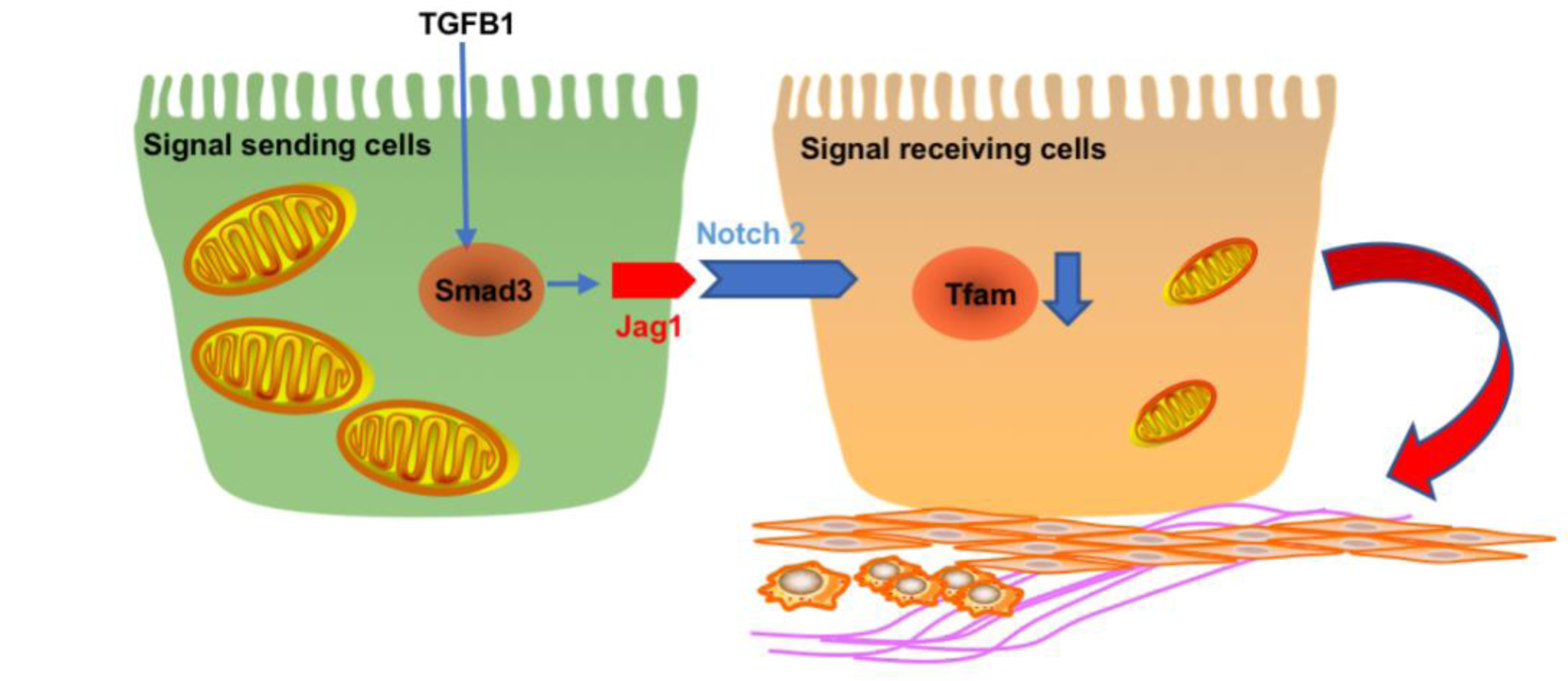
Tubule specific *Jag1/Notch2* signaling plays a key role in kidney fibrosis development by regulating metabolism via *Tfam* expression.

## Discussion

Here we showed that the Notch ligand *Jag1* and the Notch receptor *Notch2* expression in renal tubule cells play a key role in kidney fibrosis development. While previous studies from our and other groups have already suggested the contribution of Notch signaling in chronic kidney disease, most prior studies have used models with global inhibition or activation of Notch signaling(8, 46, 47). While such studies were instrumental to the understanding of the biology of kidney fibrosis, they did not represent clinically translatable strategies for conditions such as fibrosis, as long-term systemic inhibition of Notch signaling are associated with significant side effects(18). Gamma secretase inhibitors are associated with significant intestinal toxicity as Notch signaling plays a key role in intestinal cell differentiation. To circumvent these issues, receptor and ligand isoform specific targeting are emerging with improved side effect profiles.

Here, we used mice with tubule specific genetic deletion of *Notch1*, *Notch2* and global deletion of *Notch3*, and we show that only mice with tubule specific loss of *Notch2* is protected from kidney fibrosis. Similarly, genetic removal of *Jag1* in tubule epithelial cells were also protective in animal models. These results are consistent with unbiased gene expression analysis results that indicated that the expression of *Jag1* and *Notch2* were the highest in mouse and human kidneys. The expression of *JAG1* and *NOTCH2* showed the strongest correlation with kidney fibrosis in human kidneys. In addition to our current direct genetic models, several other prior studies suggested Notch2 as an indirect target of pharmaceutical interventions for kidney fibrosis. For example, the Lorenzen group showed the miR-21 is an important upstream regulator of *Notch2* and therefore miR-21 antagonist that is currently in clinical studies might also reduce *Notch2* activation (48). Similar studies showed that Klotho deficiency can cause fibrosis via *Notch2* activation (49). It is important to note that isoform-specific Notch receptor targeting has been more difficult to develop than ligand targeting, furthermore one study examining the role of *Notch2* in podocytes in nephrotic syndrome indicated some potential benefit (50). Therefore, ligand targeting such as JAG1 appears to be the most reasonable approach for CKD. Our studies can provide the foundation for the development of novel molecular therapeutics for kidney failure and advance our understanding of the renal effect of JAG1 antagonism based therapies.

Our studies show that Notch directly binds to the metabolic transcription factor *Tfam* to reprogram tubule epithelial metabolism, and induces tubule epithelial dedifferentiation and proliferation. While prior work indicated the role of metabolic changes in kidney disease development, here we show a direct interaction between Notch and metabolic changes as *Tfam* is its direct target. Kidney tubules are highly metabolic cells and many prior studies have focused on identification of the key metabolic transcription factor for tubule cells. Studies have demonstrated a decrease in expression of *Ppargc1a*, *Ppara* and *Pparg* (51). Transgenic expression of *Ppargc1a* and *Ppara* showed protection from acute kidney injury or kidney fibrosis indicating the key role of metabolic pathways in kidney function and tubule health (44). On the other hand, mice with global deletion of *Ppargc1a* and *Ppara* did not present with renal abnormalities, thus raising significant doubts that the reduced expression of these transcription factors are actually important for fibrosis development (43, 52). Here, we show that *Tfam* is the essential transcription factor for renal tubule cell metabolism. Tubule specific deletion of *Tfam* in mice results in tubule epithelial dedifferentiation and perinatal lethality. These results indicate that while transgenic expression of *Ppargc1a* and *Ppara* showed protection, the key metabolic transcription factor in the kidney is *Tfam* as no other transcription factor was able to compensate for its loss(51).

We have also analyzed other targets. We found direct Notch binding to the *Egf* promoter and subsequent inverse correlation between Notch levels and *Egf* expression in the kidney. Our *in vitro* studies failed to establish a causal relationship between Notch-regulated *Egf* levels and subsequent kidney fibrosis development. Overall, our studies indicate that Notch-induced global changes in differentiation was mediated by the regulation of cellular metabolism rather than individual targeting individual differentiation-associated gene expression. Given the broad transcriptional program Notch needs to regulate, it actually makes sense that regulated cellular metabolism alters cell differentiation states.

Future studies shall aim to understand the mechanism of sustained and excessive Notch activation in CKD. Increased tonic Notch activation is likely important in orchestrating the regenerative response including transit amplification and proliferation. It seems, however, that in CKD, Notch activation is excessive and sustained, which might be caused by the sustained epithelial injury. Overall, our studies indicate that targeting specific Notch ligand JAG1 or NOTCH2 could have important therapeutic potential for the treatment of chronic kidney disease.

## Methods

### Mice

Mice with *Jag1*-floxed alleles were kindly provided from Dr. Verdon Taylor. Mice with a floxed *Notch1* allele were purchased from Jackson Lab (Stock#006951). Mice with *Notch2* floxed allele were a gift from Dr. Ursula Zimber-Strobl. *Tfam^flox/flox^* mice were purchased from Jackson Lab (Stock#026123). *Jag1^flox/flox^, Notch1^flox/flox^, Notch2^flox/flox^,* and *Tfam^flox/flox^* mice were crossed with transgenic mice expressing Cre recombinase under the cadherin 16 promoter (*Ksp-Cre*) (Jackson Lab, stock#012237). *Notch3-/-* mice were purchased from Jackson Lab (Stock#010547). 6-8 weeks old male mice were injected with 250mg/kg FA in 300mM sodium bicarb and sacrificed on day 7 post.

### Gene expression analysis

RNA sequencing data from FA treatment for 3d mouse kidney samples were compared with normal group (GSE65267, GSM1591198-GSM1591199 vs GSM1591206-GSM1591208)(36). UUO 8d mice were compared with their sham operated group (GSE79443, GSM2095449-GSM2095452 vs GSM2095456-GSM2095458)(37). Mice with transgenic expression of APOL1 risk variants in podocytes were compared with control littermates (GSE81492, GSM2154804-GSM2154809 vs GSM2154800-GSM2154803) (35).

### Human samples

95 tubule compartments of human kidney samples were microdissected from nephrectomies as previously described (ArrayExpress: E-MTAB-5929, E-MTAB-2502) (38). Fibrosis scores for all samples were quantified by a pathologist using PAS-stained kidney sections. Correlation between fibrosis score and gene expression used a linear regression model adjusted for gender, age, and race.

### RNA sequencing

Primary culture of mouse renal epithelial cell and TGF-β1 treatment was done as previously described (44). RNA quality was assessed with the Agilent Bioanalyzer 2100 and only samples with RIN scores above 7 were used for cDNA production. RNA (100 ng) was used to isolate poly(A)-purified mRNA using the Illumina TruSeq RNA Preparation Kit. Single end 50-bp sequencing was performed in an Illumina HiSeq, and the annotated RNA counts (fastq) were calculated by Illumina’s CASAVA 1.8.2. Sequence quality was surveyed with FastQC. Adaptor and lower-quality bases were trimmed with TrimGalore. Reads were aligned to the Gencode mouse genome (GRCm38) using STAR-2.4.1d. Read counts for each sample were obtained using HTSeq-0.6.1 (htseqcount).

### Staining

Kidneys were harvested from control and FA injected mice or human kidneys. Histological analysis was examined on formalin-fixed, paraffin-embedded kidney sections stained by PAS (periodic acid Schiff). Immunofluorescence staining was performed with antibodies against JAG1 (Santa Cruz#SC9303) and NOTCH2 (Millipore#07-1234) in OCT-embedded frozen kidney sections. Tubule specific markers were used as previously described (53). Immunohistochemistry staining was performed with antibody against COX IV (Abcam# ab16056) in paraffin-embedded kidney sections.

### BUN and Creatinine level

Serum creatinine and BUN were determined by Mouse Creatinine Kit (Crystal Chem) and TRACE DMA Urea kit (Thermo Electron Corporation), respectively, according to the manufacturers’ instructions.

### qRT-PCR

RNA was isolated from harvested kidney tissues and cells using Trizol (Invitrogen). RNA (1 μg) was reverse transcribed using the cDNA Archival Kit (Life Technologies), and qRT-PCR was run in the ViiA 7 System (Life Technologies) instrument using SYBR Green Master Mix and gene-specific primers. The data were normalized and analyzed using the ΔΔCt method. The primers used are listed in **Supplemental Table 3**.

### Cell culture

Rat epithelial cells (NRK52E) from ATCC (CRL-1571) were cultured in DMEM with 5% FBS. L cell and JAG1-expressing L cells were a gift from Dr. Pamela Stanley from Albert Einstein College of Medicine. For co-culture system, NRK-52E cells were firstly plated, then incubated overnight and reached 70%-80% confluent the next day. Then, L cell and JAG1-expressing L cells were plated on top of signal-receiving cells at a density equal to the monolayer to establish a direct contact between ligand and receptor. The cells were co-cultured in growth medium for 24 hours. For the rescue experiment, NRK-52E were either transiently co-transfected with *TFAM*-expressing plasmid and GFP as control or EGF. Transfection efficiency was confirmed by GFP visualization. Following transfection, the cells were co-cultured with either JAG1-expressing L cells or L cells for 24 hours, or were treated with TGF-β1 (5 ng/ml) for 48 hours. pCellFree_G03 *TFAM* was a gift from Dr. Kirill Alexandrov (Addgene plasmid # 67064) (54). Recombinant Human EGF was purchased from Peprotech.

### ChIP-seq

RBPJ ChIP-seq dataset with 6h and 24h Notch activation and inhibition was adapted from cultured cells (GSE37184) (41).

### Oxygen consumption experiment

FAO were studied using SeaHorse Bioscience metabolic analyzer as previously described (44). We followed manufacturer’s instruction to monitor oxygen consumption rate (OCR) in TGF-β1-treated NRK52E cells or co-culture system. Briefly, cells were seeded in XF24 V7 cell culture microplate at 2 × 10^4^ cells per well. OCR was assessed at baseline and after the addition of palmitate conjugated BSA (180 μM) followed by the addition of the carnitine palmitoyltransferase-1 inhibitor etomoxir (40 μM). The final state was determined after the addition of the ATP synthase inhibitor oligomycin (1 μM).

### ATP measurement

The ATP concentration of cultured cells was measured using an ATP Fluorometric Assay Kit (BioVision) according to the manufacturer’s protocol.

### Statistics

Statistical analyses were performed using GraphPad Prism 6.0 software. All values are expressed as mean and standard deviation. A 2-tailed Student’s *t* test was used to compare 2 groups. One-way ANOVA or Two-way ANOVA with *post*-*hoc* Tukey test were used to compare multiple groups. A *P* value less than 0.05 was considered statistically significant.

### Study approval

All experiments in animals were reviewed and approved by the Institutional Animal Care and Use Committee of the University of Pennsylvania, and were performed in accordance with the institutional guidelines.

## Author contributions

S.H. and K.S. designed research studies. S.H., Y.S., S.Y.H., and S.Y.L conducted experiments. S.H., J.P., and C.Q. analyzed data. U.Z-S. and V.T. provided transgenic mice. S.H. and K.S. wrote the manuscript.

## Acknowledgments

The authors thank Dr. Pamela Stanley from Albert Einstein College of Medicine for providing L cell and JAG1-expressing L cells. The authors thank Frank Chinga and Antje Gruenwald for assistance with animal breeding and care. The authors thank Joshua Bryer for proofreading. The authors thank the Molecular Pathology and Imaging Core at University of Pennsylvania for histology services. This work was supported by the NIH R01 DK076077, DK087635, and DK1088220 and by the Juvenile Diabetes Research Foundation grants to K.S.

